# Effect of Changing Heart Rate on the Ocular Pulse and Dynamic Biomechanical Behavior of the Optic Nerve Head

**DOI:** 10.1101/770917

**Authors:** Yuejiao Jin, Xiaofei Wang, Sylvi Febriana Rachmawati Irnadiastputri, Rosmin Elsa Mohan, Tin Aung, Shamira A. Perera, Craig Boote, Jost B. Jonas, Leopold Schmetterer, Michaël J.A. Girard

## Abstract

**Purpose:** To study the effect of changing heart rate on the ocular pulse and the dynamic biomechanical behaviour of the optic nerve head (ONH) using a comprehensive mathematical model.

**Methods:** In a finite element model of a healthy eye, a biphasic choroid consisted of a solid phase with connective tissues and a fluid phase with blood, and the lamina cribrosa (LC) was viscoelastic as characterized by a stress-relaxation test. We applied arterial pressures at 18 ocular entry sites (posterior ciliary arteries) and venous pressures at four exit sites (vortex veins). In the model, the heart rate was varied from 60 bpm to 120 bpm (increment: 20 bpm). We assessed the ocular pulse amplitude (OPA), pulse volume, ONH deformations and the dynamic modulus of the LC at different heart rates.

**Results:** With an increasing heart rate, the OPA decreased by 0.04 mmHg for every 10 bpm increase in heart rate. The ocular pulse volume decreased linearly by 0.13 µL for every 10 bpm increase in heart rate. The storage modulus and the loss modulus of the LC increased by 0.014 MPa and 0.04 MPa, respectively, for every 10 bpm increase in heart rate.

**conclusions:** In our model, the OPA, pulse volume, and ONH deformations decreased with an increasing heart rate, while the LC became stiffer. The effects of blood pressure / heart rate changes on ONH stiffening may be of interest for glaucoma pathology.

**Support:** Singapore Ministry of Education, Academic Research Fund, Tier 2 (R-397-000-280-112).

**Commercial relationship:** None

## INTRODUCTION

Glaucoma is a complex, multifactorial disease. Although the pathogenic mechanism of glaucoma has still not been well understood, it has generally been accepted that various interlinked biomechanical and vascular pathways to glaucomatous injury are involved, and that these are closely related to the optic nerve head (ONH) biomechanics.^1-8^ The ONH, and in particular the lamina cribrosa (LC), is a primary site of interest during the development of glaucoma since it is the region where retinal ganglion cell (RGC) axon damage occurs. Because the ONH is constantly exposed to several loads in vivo (e.g. IOP, cerebrospinal fluid pressure (CSFP), orbital fat pressure, optic nerve traction),^6, 9-13^ there is a general consensus that knowing how ‘robust’ a ONH is could be a critical information for estimating the risk of developing glaucoma.

To date, to assess the biomechanics of the ONH in vivo, the typical approach has been to artificially manipulate IOP (e.g. with ophthalmodynamometry) while observing the resulting deformations with optical coherence tomography (OCT).^14-17^ While such methods may have value from a research point of view, they may not be easily translated clinically due to discomfort during the examination of the patients. However, IOP is not constant but rather pulsatile, fluctuating constantly with the cardiac cycle. The fluctuation of IOP with the heart rate has been known as the ocular pulse and the difference between the systolic and diastolic IOP has been termed the ocular pulse amplitude (OPA). In a previous study, we proposed a framework to assess ONH biomechanics in vivo without artificially manipulating IOP. ONH deformations can indeed be measured by using the ocular pulse as the load,^18^ and from such data, modelling can help us interpret these deformation data to derive stiffness estimates for the ONH. ^19^

This is however a complex engineering problem: the ONH tissues are viscoelastic,^7, 20, 21^ which basically means that ONH stiffness is affected by the rate of loading, or in other words, the duration of the cardiac cycle. Indeed, collagenous soft tissues typically appear stiffer when they are loaded faster. This is a protective mechanism for soft tissues to handle highly dynamic events.^7, 20^ Furthermore, it has been shown clinically, that an increase in heart rate or a change in blood pressure can also affect the OPA;^22, 23^ this will in turn affect the deformations of the ONH.

To measure the stiffness of the ONH using the ocular pulse as a natural load, we believe it is critical to first build an understanding of how the heart rate and blood pressure change the IOP (i.e. the OPA), and how they also affect the stiffness and deformations of the ONH tissues. From a clinical point of view, if ONH tissues appear stiffer, a physician should be able to tell whether this is because of a different heart rate / blood pressure, or because of tissue remodelling from glaucoma. In this study, we aim to build such an understanding using finite element (FE) modelling. In the future, our proposed models could be combined with experimental data to assess ONH stiffness in vivo.

## METHODS

Using FE modelling, we previously modelled the origin of the ocular pulse and its effect on the ONH.^19^ We found that a change in arterial pressure (as that occurring from diastole to systole) resulted in choroidal swelling, which in turn induced a change in IOP (the OPA) due to the incompressibility of the vitreous. Both the OPA and choroidal swelling were responsible for deforming the ONH tissues, and a relatively large shearing of neural tissues was observed within the neuroretinal rim. The vast majority of the trends we reported matched clinical and experimental data.

In this study, we used the same model, but we further defined the LC as a viscoelastic material. Such viscoelastic properties were obtained experimentally using uniaxial stress relaxation tests. Using our FE model, we then varied the heart rate from 60 bpm to 120 bpm and studied its effect on: 1) the OPA, 2) the choroidal pulse volume, 3) ONH deformations, and 4) LC stiffness.

### Three-Dimensional Geometry of the Ocular and Orbital Tissues

The geometry of the three-dimensional eye model was adapted from our previous studies.^12, 19, 24^ In summary, the eye globe and optic nerve were reconstructed from magnetic resonance imaging images of a heathy subject. The corneoscleral shell was assumed to be spherical with an outer diameter of 24 mm and thickness of 1 mm. The optic nerve was composed of three tissues: the nerve tissue, the pia mater and the dura mater. A generic ONH geometry was constructed and embedded within the corneoscleral shell, including the scleral flange, Bruch’s membrane, the LC, the prelaminar tissue and the border tissue of Elschnig and Jacoby as extensions of the pia mater. The model was then meshed using 67,584 eight-node hexahedrons and 3,024 six-node pentahedrons (Figure 1). The mesh density was numerically validated through a convergence test. The whole eye was reconstructed, and symmetry conditions were not applied.

**Figure 1.**
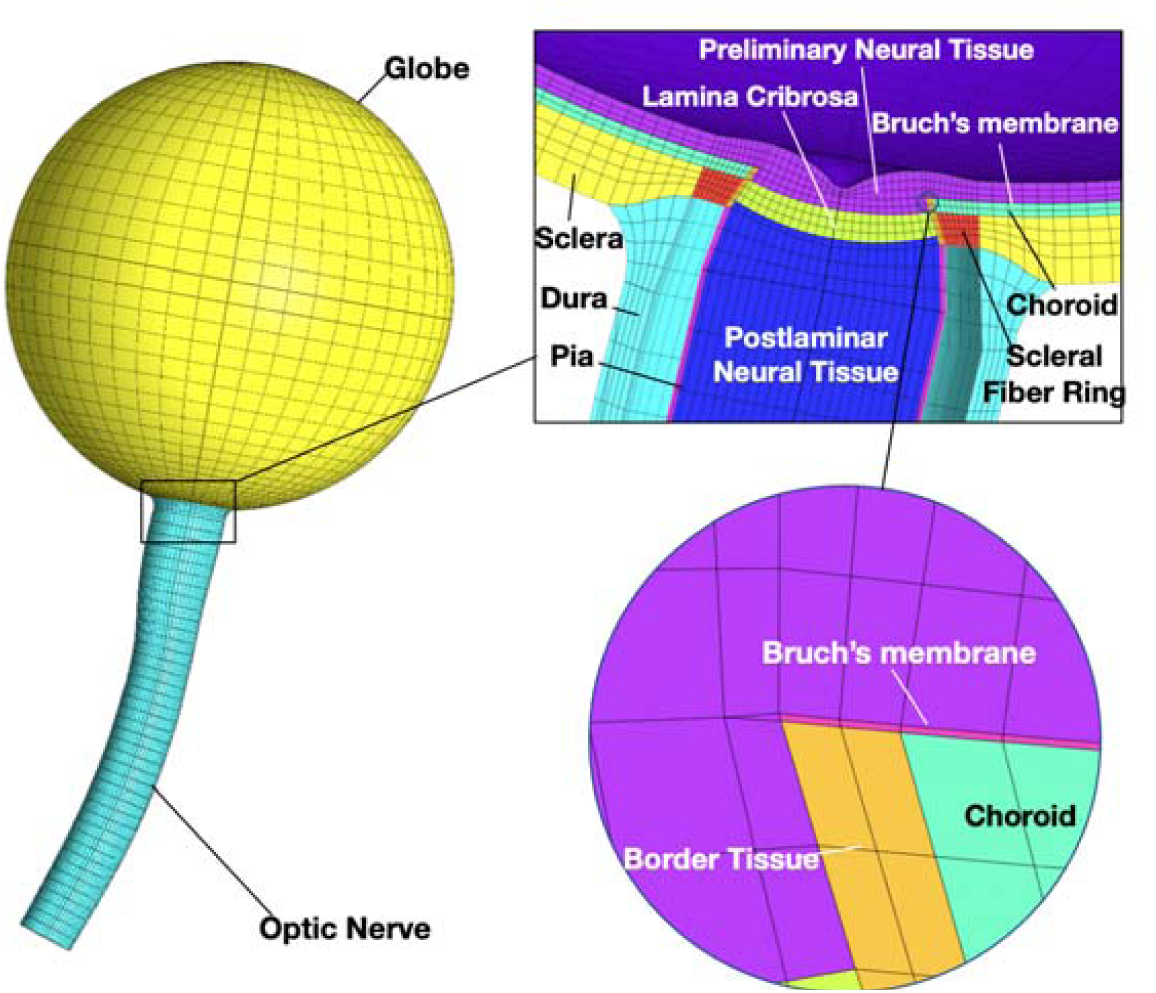
Reconstructed geometry and FE mesh of the whole eye model adopted from Jin.^19^ ONH structures (including the LC, sclera, neural tissues, choroid, BM, border tissue, pia and dura) were reconstructed using average measurements from the literature.^9, 19, 24, 26, 68^

### Biomechanical Properties of the Ocular Tissues

The sclera was modelled as a fiber-reinforced composite, as described in our previous paper.^25^ The neural tissues and Bruch’s membrane were modelled as isotropic elastic materials. ^26^ The pia and dura were modelled as Yeoh materials, as characterized from experiments with porcine eyes.^24^ As performed in our previous study, we modelled the choroid as a biphasic material comprising a solid phase and a fluid phase (i.e. blood) to allow for changes in blood pressure during the cardiac cycle and the movement of blood within the choroid.^19^ All biomechanical parameters are listed in **Table 1**.

**Table 1.** Tissue biomechanical properties used for all models.

| Tissue | Constitutive Model | Biomechanical Properties | References |
| --- | --- | --- | --- |
| Sclera | Mooney-Rivlin Von Mises<br>distributed fibers | $c_1 = 0.285 \text{ MPa}$<br>$c_3 = 0.0137 \text{ MPa}$<br>$c_4 = 658.125$<br>$k_f = 2$ (scleral ring)<br>$k_f = 0$ (other region of sclera)<br>$\theta_p$: preferred fiber orientation | Girard et al.<sup>25</sup> |
| LC | Viscoelastic | Instantaneous modulus = 8.49 MPa<br>Equilibrium modulus = 0.49 MPa<br>Long-term time constant = 272 s<br>Short-term time constant = 8.33 s | Porcine tissue experiment (this study) |
| Neural tissue | Isotropic elastic | Elastic modulus = 0.03 MPa<br>Poisson's ratio = 0.49 | Miller<sup>69</sup> |
| BM | Isotropic elastic | Elastic modulus = 10.79 MPa<br>Poisson's ratio = 0.49 | Chan et al.<sup>70</sup> |
| Choroid | Biphasic | solid volume fraction: 0.55<br>Solid matrix:<br>Neo-Hookean<br>Elastic modulus = 0.06 MPa<br>Poisson's ratio = 0<br>Permeability: 45037 mm<sup>2</sup>/MPa.s | Jin et al.<sup>19</sup> |
| Dura | Yeoh model | $C_1 = 0.1707 \text{ MPa}$<br>$C_2 = 4.2109 \text{ MPa}$<br>$C_3 = -4.9742 \text{ MPa}$ | Wang et al.<sup>24</sup> |
| Pia | Yeoh model | $C_1 = 0.1707 \text{ MPa}$<br>$C_2 = 4.2109 \text{ MPa}$<br>$C_3 = -4.9742 \text{ MPa}$ | Wang et al.<sup>24</sup> |
| Border Tissue | Isotropic elastic | Elastic modulus = 10.79MPa<br>Poisson's ratio = 0.49 | Chan et al.<sup>70</sup> |

### Viscoelastic Properties of the Lamina Cribrosa

In order to characterize the response of the LC under dynamic loading such as the heart rate, we studied its viscoelastic properties. Since the viscoelasticity properties of the LC have not been reported in the literature yet, we conducted uniaxial stress relaxation tests of the LC of seven porcine eyes.

Experiments were performed in accordance to the ARVO (Association for Research in Vision and Ophthalmology) statement for the Use of Animals in Ophthalmic and Vision Research. The experimental procedure followed the previously described methodology designed for the sclera.^27, 28^ In brief, seven fresh porcine eyes were collected and cleaned of all extraorbital tissues. After dissecting the eye and cutting the optic nerve, the retina and choroidal layers were removed. A customized device was used to cut a scleral strip (orientation: inferior to superior) with the LC in its center (width: 1.2 mm; Figure 2). Mineral oil was applied to the specimen surface to maintain hydration. Each sample was mounted between the two grips of the uniaxial tensile tester (Instron 5848; Instron. Inc., Noor wood, MA, USA) with the LC exposed in the center (Figure 2c). The sample was subjected to 10 cycles of preconditioning at 1% strain prior to the stress relaxation test. The sample was then subjected to a maximum of 5% strain at a rate of 1% per second, followed by a relaxation period of 300s at constant stress. These strain levels were chosen as they are typically exhibited in vivo under physiological conditions (during eye movements or change in IOP).^10^ The thickness measurements of the LC (mean: 0.628mm) were obtained by spectral-domain optical coherence tomography (Spectralis OCT, Heidelberg Engineering, Heidelberg, Germany) and were used to compute the tensile stress. The averaged stress values of seven porcine eyes were fitted with a linear viscoelastic model incorporating a reduced relaxation function to characterize the viscoelastic properties of the LC.^28^ A typical stress relaxation curve is shown in Figure 2d.

**Figure 2.**
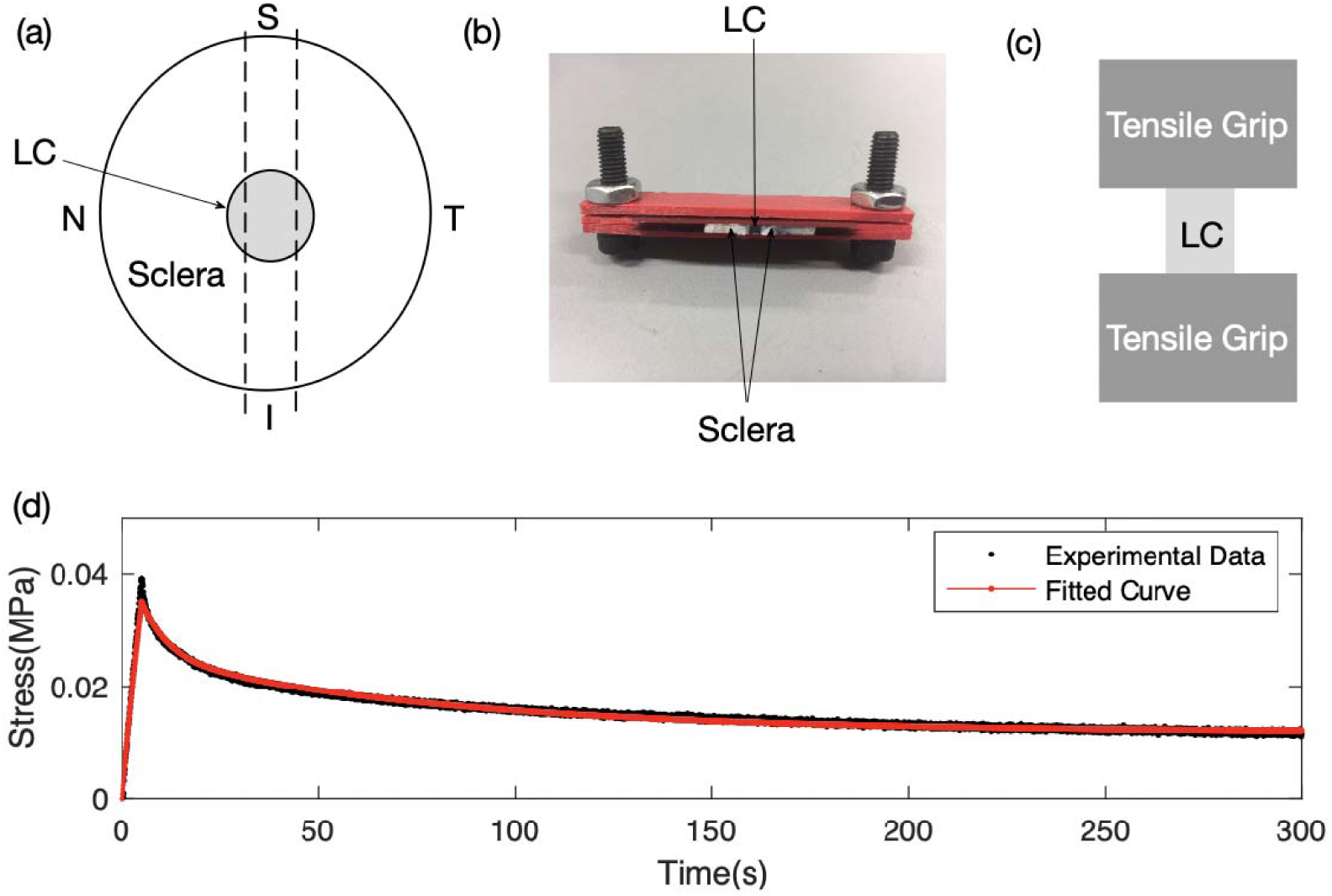
(a) Schematic showing the scleral strip with the LC in its center. The uniaxial tensile specimen was obtained by using a customized device (shown in b) to obtain a fixed width of 1.2 mm. It was then clamped between the tensile grips (shown in c) with the LC exposed in its centre. (d) A typical stress relaxation experimental test dataset for an LC sample fitted with a linear viscoelastic model curve.^28^

### Boundary and Loading conditions

Two sets of boundary conditions were applied to ensure numerical stability. First, the orbital apex of the optic nerve was fixed to model its connection to the optic canal. Second, the eye globe was fixed near the equator on two opposite sides with four nodes on each side. This was carried out to improve stability and to avoid such boundary conditions from having an impact on the ONH deformations.

We applied a baseline IOP of 15 mmHg to the inner limiting membrane and a baseline CSFP of 11.3 mmHg to the arachnoid space (Figure 3).^29^ As described previously, we applied prescribed fluid pressures on the choroidal layer to mimic the blood flow inlets (long and short posterior ciliary arteries [PCAs]) and outlets (vortex veins).^19^ The arterial pressure was varied between 70.8 and 93 mmHg to mimic the diastolic and systolic changes of the ophthalmic artery pressure.^30^ A constant blood pressure of 15 mmHg was applied at the four vortex veins exits out of the choroid.^31^ In our model, it was the pressure difference between the PCAs and vortex veins that drove the choroidal blood flow, allowing the choroid to swell during the cardiac cycle.

**Figure 3:**
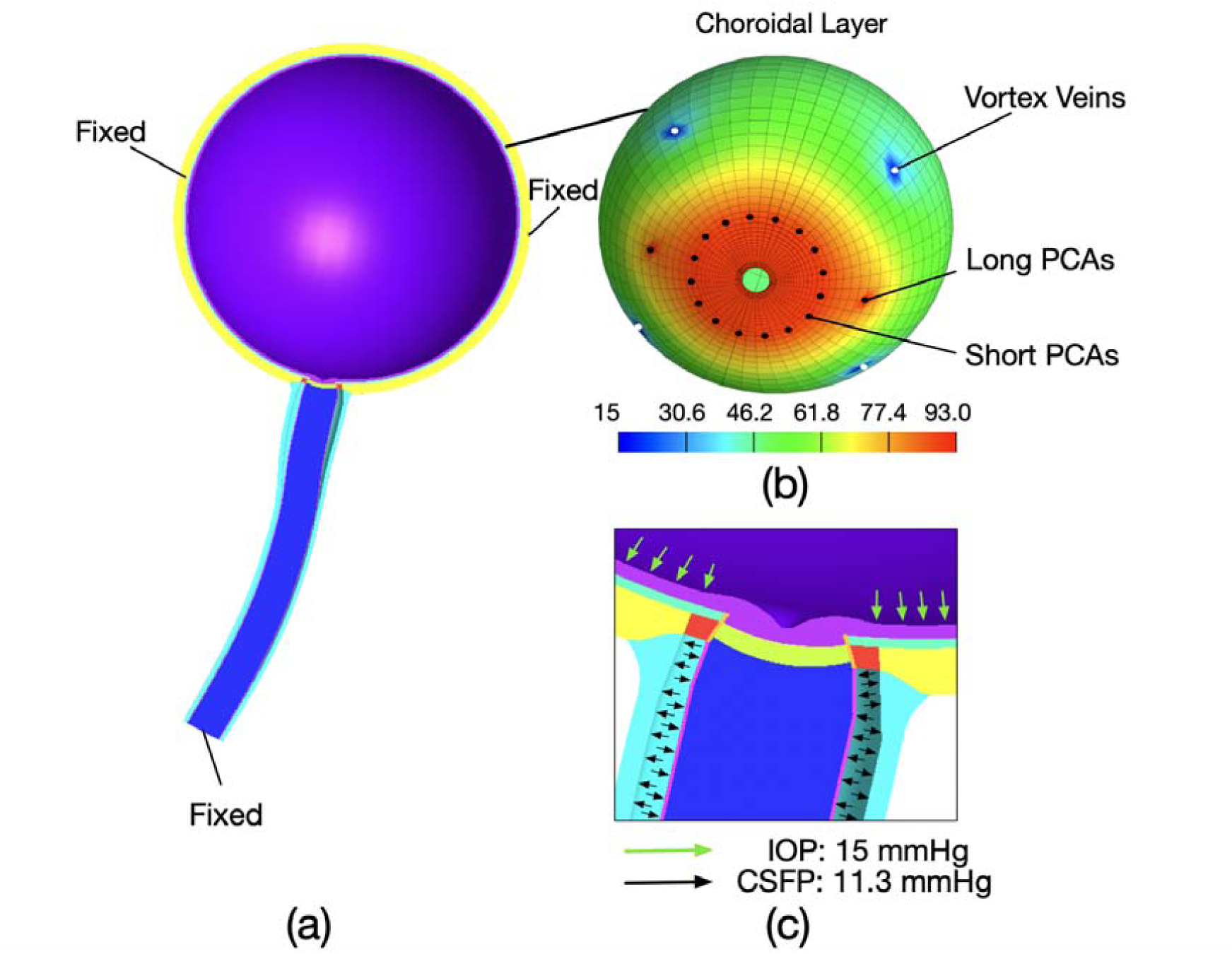
(a) Half eye geometry and boundary conditions: fixed optic nerve at the orbital apex and fixed sclera nodes near the equator on two opposite sides. (b) Choroidal with the nodes where arterial blood pressure (short and long PCAs) and venous blood pressures (vortex veins) were applied. (c) ONH region illustrating the pressure loads (IOP, CSFP) applied in each model.

### Changing the Heart Rate from 60 to 120 bpm

We aimed to understand the impact of different heart rates on the ocular pulse, ONH deformations, and LC stiffness. Hence, four models were implemented with the heart rate ranging from 60 to 120 bpm with an increment of 20 bpm. All the boundary conditions and material properties were kept constant, such that only the heart rate varied.

### FE Processing to Predict the OPA, Pulse Volume, and ONH Deformations

All FE models were solved with FEBIO v2.8.5 (Musculoskeletal Research laboratories, University of Utah, Salt Lake City, UT, USA).^32^ The OPA was estimated by first assuming that the vitreous was incompressible as performed in our previous study.^19^ This is a reasonable assumption because the vitreous mostly consists of water, which is also incompressible, and is gas-free. Therefore, the volume of the vitreous body was constrained to remain constant during choroidal swelling in all our simulations. Since our model aimed to reproduce choroidal swelling during the cardiac cycle, such a swelling will try to deform and change the volume of the vitreous body. Since the vitreous body was constrained as incompressible, an internal pressure needed to be applied to maintain its volume. This pressure term is an output of our model and can be understood as the OPA.

More specifically, each model was implemented in three steps as shown in Figure 4. The initial step applied boundary conditions and loads including baseline IOP, CSFP and ophthalmic arterial pressure and vortex vein pressure. During the second step, a 5000s relaxation period was applied due to the viscoelasticity of the LC. All the boundary conditions and loads remained unchanged. During the final step, we imposed a volume constraint to the inner limiting membrane to ensure the incompressibility of the vitreous.^19^ The CSFP, baseline IOP and vortex vein pressure were kept constant; the ophthalmic arterial pressure varied from diastole to systole periodically following a sinusoidal profile for two cycles.

**Figure 4.**
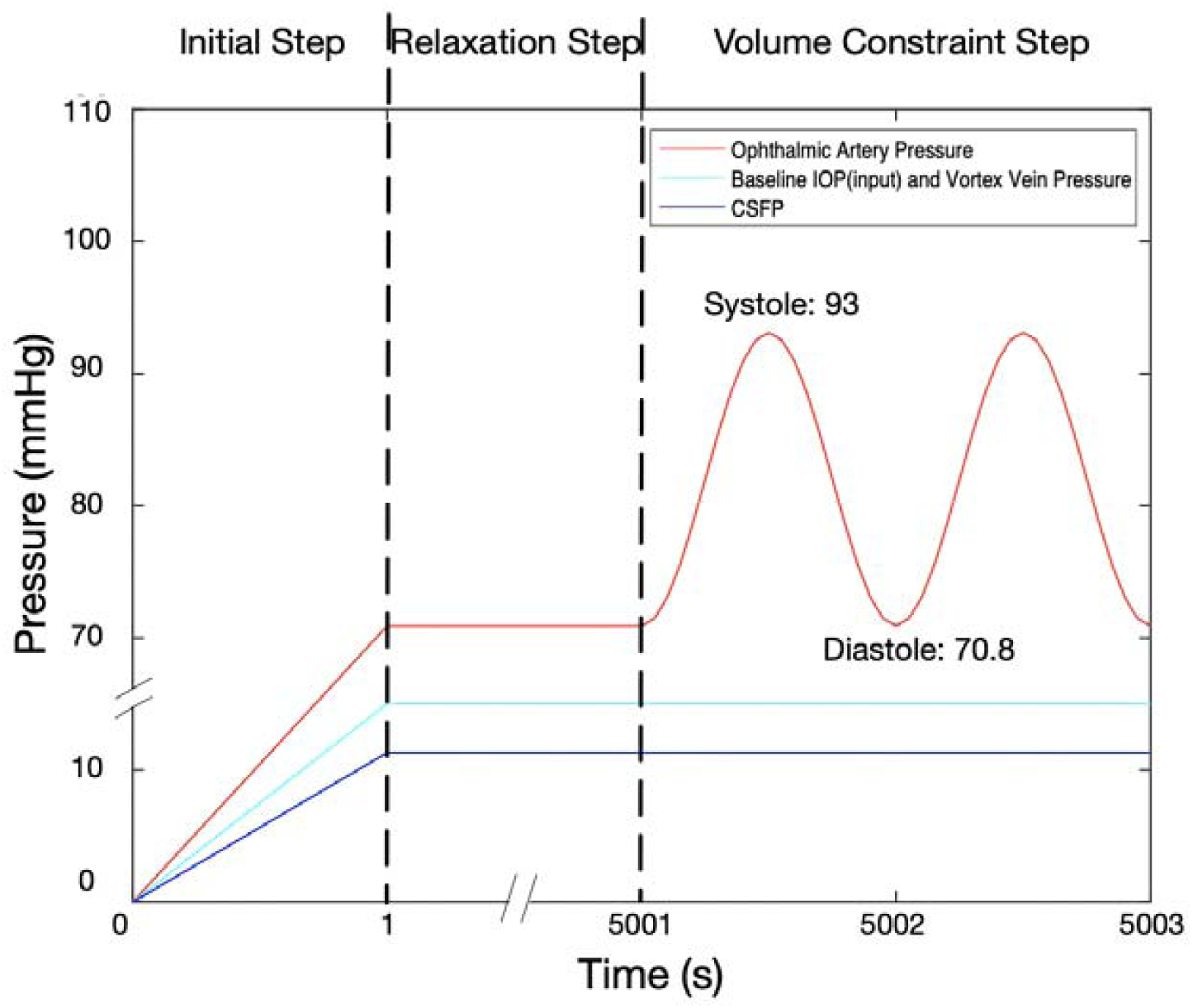
Profiles of different pressure loads.

The post processing was performed using MATLAB (v2018; Mathworks, Natick, MA, USA). For each model, we reported the OPA, the pulse volume, the diastole-to-systole displacements of the LC and of the pre-lamina tissue. In addition, we characterized the stiffness of the LC at different heart rates in the central region along the radial, circumferential and longitudinal directions (Figure 5).

**Figure 5:**
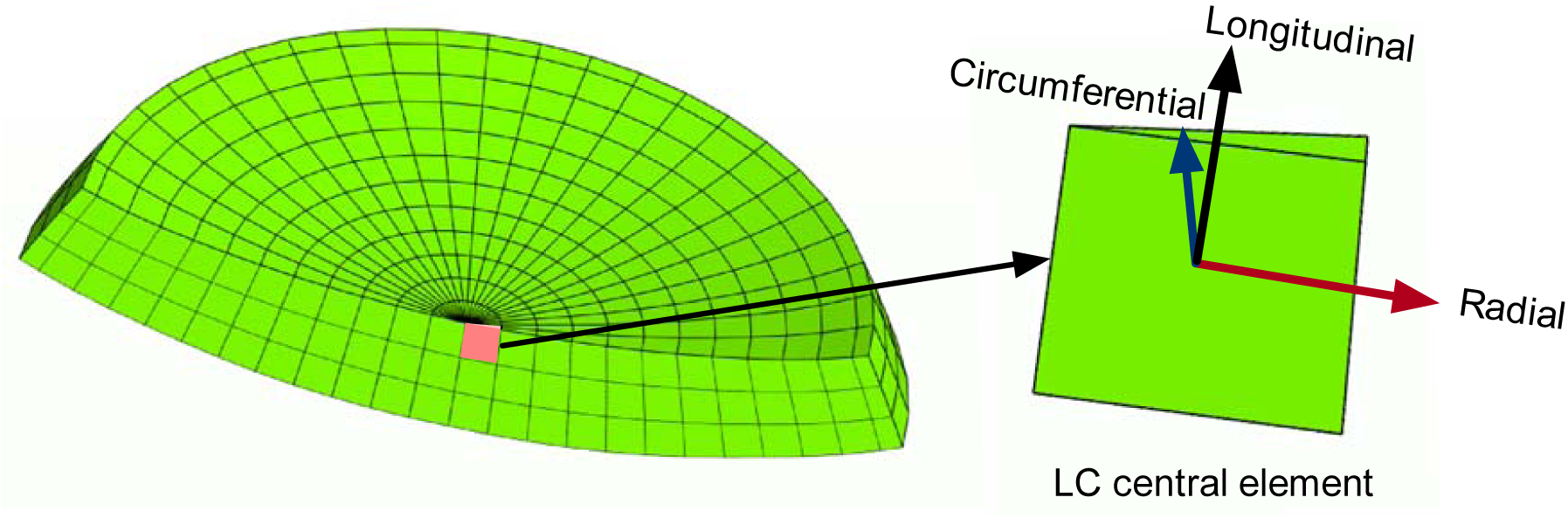
Cross-section of the LC illustrating the local coordinates (radial, circumferential and longitudinal) for one central element.

### Characterizing LC Stiffness with Changing Heart Rate

The stiffness of the LC can be characterized by computing its dynamic modulus, as typically performed for dynamic mechanical analyses.^20^ Since the ocular pulse induced a cyclic deformation of the LC, the LC strain ‘input’ can be represented as:

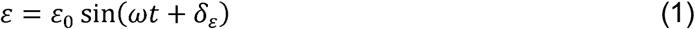

where *ε*_0_, *δ*_*ε*_, and *ω* are the peak strain amplitude, strain phase shift and angular frequency of the harmonic excitation, respectively. Since the LC only exhibits small strains, the LC can be approximated to be linear viscoelastic.^33^ Hence, the resulting sinusoidal stress ‘output’ would differ in phase from the ‘input’ strain depending on the viscoelastic properties of the LC, and can be expressed as:

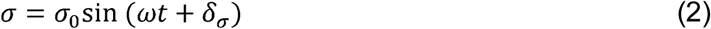

where *σ*_0_ and *δ*_*σ*_are the peak stress amplitude and stress phase shift, respectively. The phase delay between the stress and the strain due to viscoelastic effects is known as the phase lag *δ*, i.e. the difference between *δ*_*ε*_and *δ*_*σ*_(Figure 6). The dynamic mechanical properties of the LC can now be represented by the complex dynamic modulus, consisting of the storage modulus (elastic component) and the loss modulus (viscous component). It is defined as:

**Figure 6.**
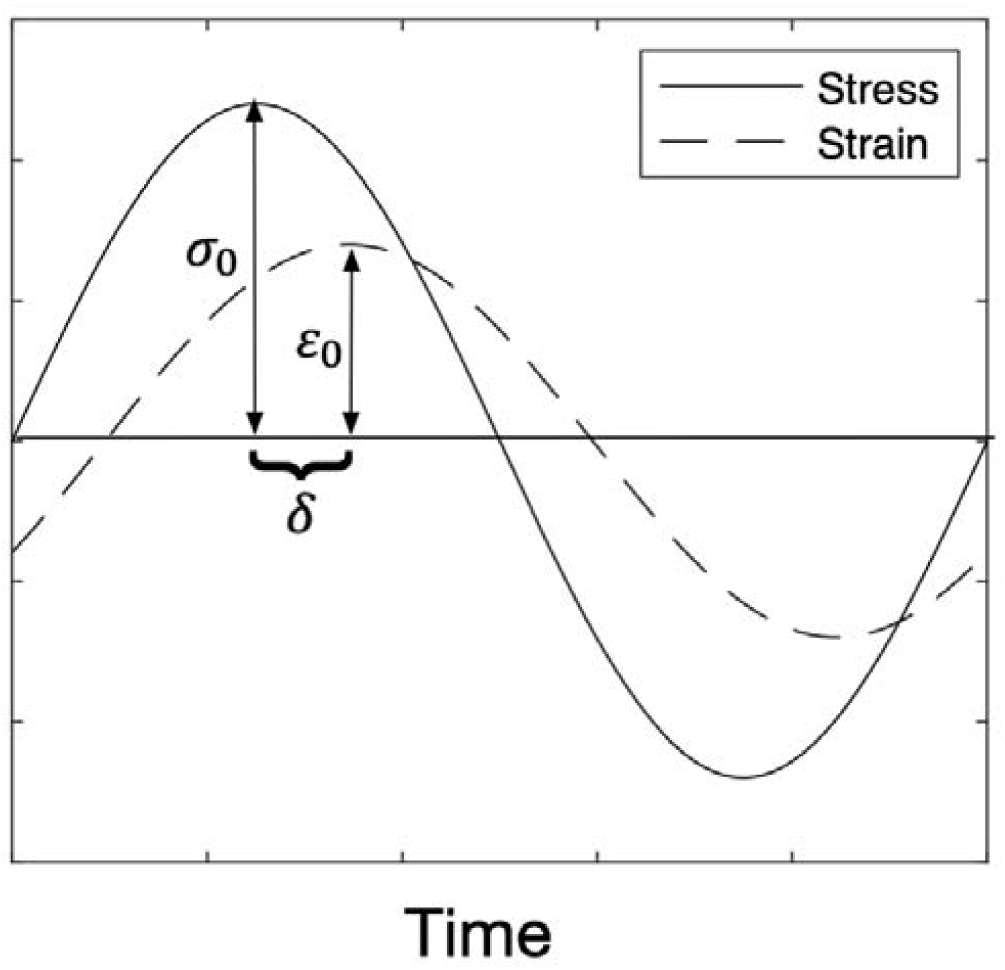
Schematic representation of the strain and stress profiles as a function of time in calculating the dynamic properties of the LC.

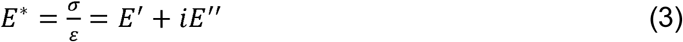

where *E*′ is the storage modulus, *E*′′ is the loss modulus, and *i* the unit imaginary number. The storage and loss moduli are related to the phase lag through the following equations:

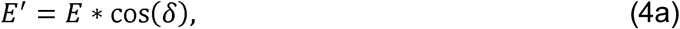

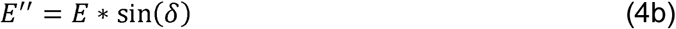

For each heart rate, we computed the storage and loss moduli in the central region of the LC (composed of one element). We first computed the Lagrange strain and Cauchy stress tensors, and estimated the diastole-to-systole Lagrange strain and Cauchy stress profiles along three directions (radial, circumferential, and longitudinal). Each strain/stress profile was then fitted with equations (1-2) to determine the peak strain/stress amplitudes and the phase shifts. From these data we extracted the phase lag, and thus the storage and loss moduli of the LC central region along 3 directions: radial, circumferential, and longitudinal.

## RESULTS

### Effect of the Heart Rate on the Ocular Pulse

For each model, the pulsating arterial pressure during the cardiac cycle resulted in choroidal swelling (increase in pulse volume), which in turn increased the IOP (the delta change in IOP being the OPA). The IOP and the pulse volume as functions of time are shown in Figure 7 for two cycles at a heart rate of 60 bpm.

**Figure 7.**
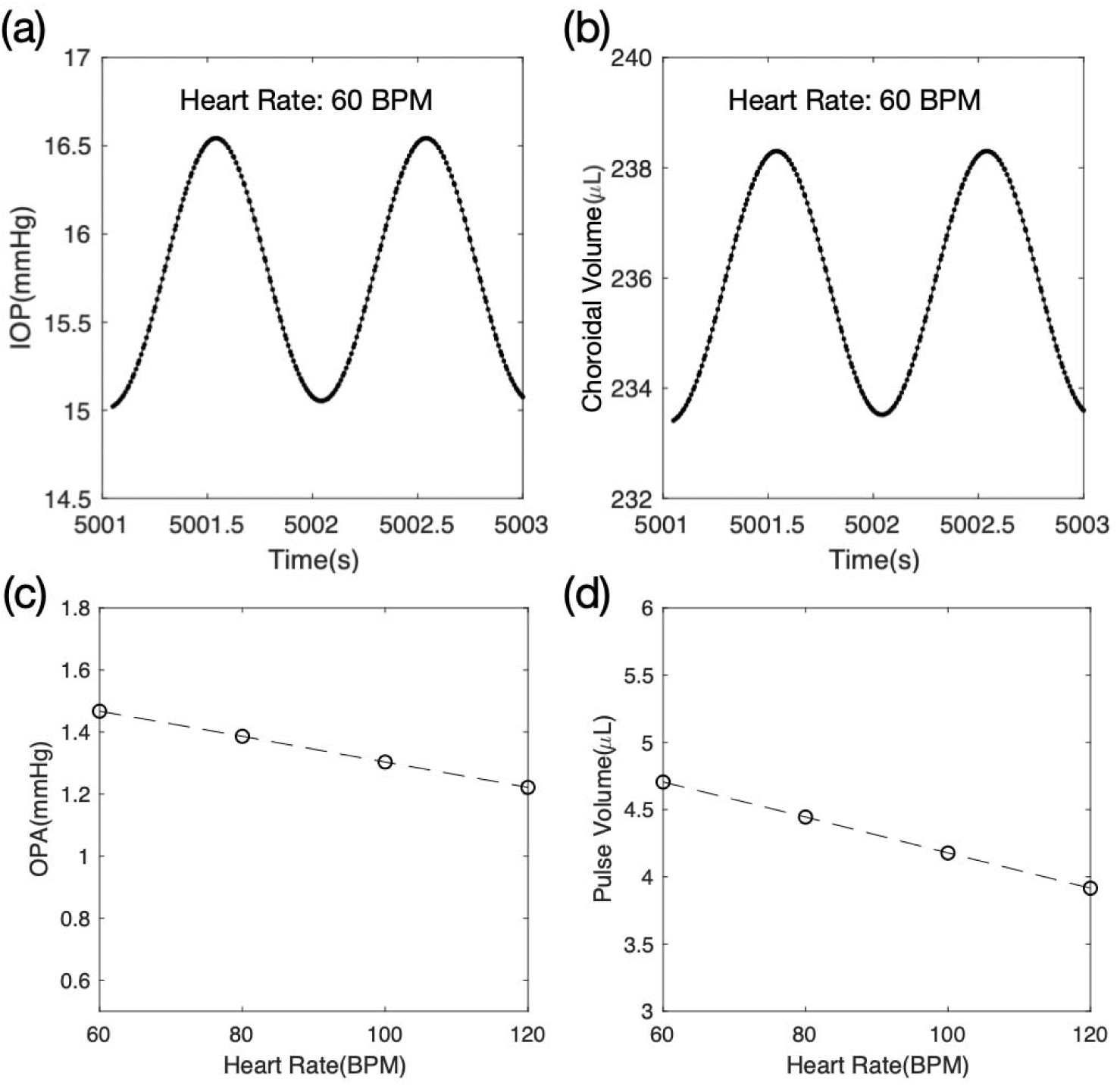
(a,b) IOP and choroidal volume over two cycles at a heart rate of 60 BPM. (c,d) Change of the OPA and the pulse volume (PV) with heart rate.

We also found that both the OPA and the pulse volume decreased with an increasing heart rate (Figure 7). On average, the OPA and the pulse volume decreased by 0.04 mmHg (2.7%) and 0.13µL (2.8%), respectively for every 10 bpm heart rate increment.

### Effect of the Heart Rate on ONH Deformations

During the cardiac cycle, the ONH moved posteriorly (diastole-to-systole) and anteriorly (systole-to-diastole). The diastole-to-systole displacement of the central anterior LC point was 5.8 µm at a heart rate of 60 bpm and decreased by 0.1 µm for every 10 bpm heart rate increment (Figure 8a-c).

**Figure 8.**
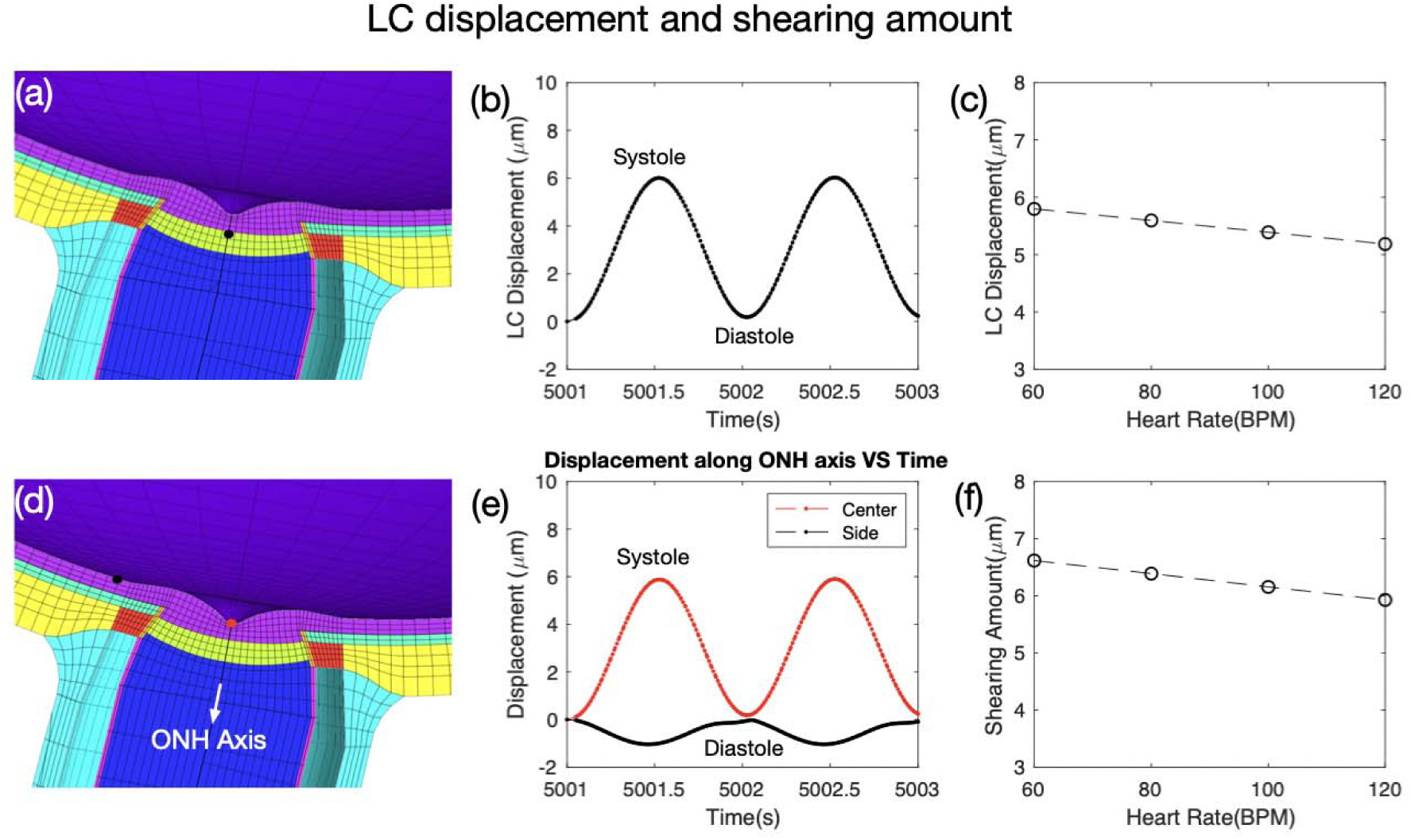
(a) Cross-section of the ONH highlighting the center of the LC and (b) the displacement of the LC center. (c) Change of the LC displacement with increasing heart rate. (d,e) Cross-section of the ONH and the radial displacement of the highlighted points (red: center of prelaminar; black: a point on the retinal surface located 0.65 mm away from BMO) over two cardiac cycles. (f) Change of the shearing amount with heart rate.

**Figure 9.**
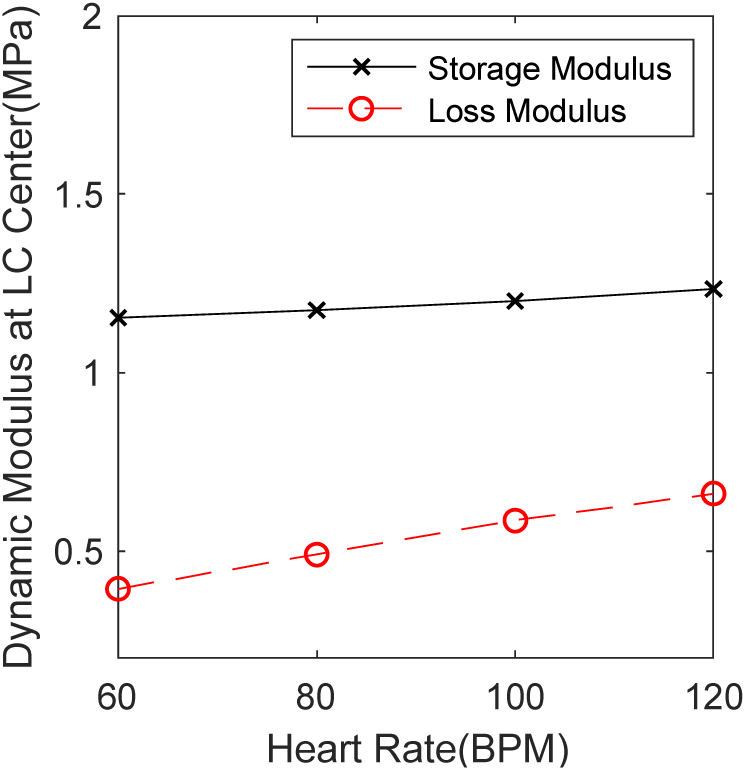
Change of the storage and loss moduli of the LC center with the heart rate.

We also observed that the prelamina (neural tissue anterior to the lamina) and the parapapillary retina moved along opposite directions during the cardiac cycle. Specifically, from diastole to systole, the prelamina moved posteriorly (as highlighted in red in Figure 8d), whereas the parapapillary retina (highlighted in black in Figure 8d) moved anteriorly. This resulted in a net shearing of the neural tissues in the neuro-retinal rim. This shearing amount (difference between the prelamina displacement and the parapapillary retina displacement) also decreased with an increasing heart rate. On average, the shearing amount reduced by 0.12 µm for every 10 bpm heart rate increment (Figure 8f).

### Effect of the Heart Rate on LC stiffness

The storage and loss moduli were computed in the central region of the LC for 3 orientations (radial, circumferential, and longitudinal). They all followed the exact same trend and all increased with an increasing heart rate. On average, the storage modulus increased by 0.014 MPa, and the loss modulus by 0.04 MPa, for every 10 bpm heart rate increment. This indicates that the lamina cribrosa will appear stiffer with a higher heart rate.

## DISCUSSION

In this study, we used FE modelling to study the impact of the heart rate on the ocular pulse and its biomechanical impact on the ONH. Our model predicted that the OPA, pulse volume and ONH deformations decreased with an increasing heart rate. In addition, the LC became stiffer at faster heart rates. These findings may help develop technologies to finely assess the biomechanical properties of the ONH in vivo.

### The OPA and the Pulse Volume Decreased with an Increasing Heart Rate

With an increase in heart rate, both the OPA and the pulse volume reduced when simultaneously the ophthalmic arterial and venous vein pressures were kept constant. A faster heart rate is associated with a shorter duration for each cardiac cycle, suggesting that the volume of blood being pushed into the choroid would be reduced for each pulse. Our results are also consistent with a previous clinical study,^23^ in which all the subjects were fitted with cardiac pacemakers and the heart rate was altered by resetting the pacemakers. Neither IOP nor systolic-diastolic blood pressures were found to be significantly altered during the experiments, suggesting that the heart rate was one of the main contributing factor causing a reduction in OPA.

It has also been observed that a short period of dynamic exercise can also lead to a reduction in OPA and ocular pulse volume.^34-36^ However, the impact of exercise is complex as it involves changes in multiple physical parameters including not only the IOP and heart rate, but also the systolic and diastolic blood pressures, and blood distribution in the body. Although heart rate and blood pressures involve separate mechanisms, exercises would induce an increase in both heart rate and blood pressures.^37^ The systolic and diastolic blood pressures are related to the ocular pulse through the ocular perfusion pressure (OPP), defined as the difference between the mean blood pressure and IOP. In one study, there was a significant reduction in IOP, and an increase in OPP immediately after strenuous exercise (resting: heart rate= 68±7.9 bpm, IOP= 19.8±5.4 mmHg, OPP = 51.2±4.9 mmHg; after exercise: heart rate=125±24.9 bpm, IOP=14.3±5.0 mmHg, OPP=56.9±7.5 mmHg).^36^ Our models could potentially be used to reproduce more complex scenarios such as those.

### ONH deformations and Neural Tissue Shear Decreased with an Increasing Heart Rate

Our model predicted that ONH deformations decreased with an increasing heart rate. It has been shown that isometric exercise caused a significant increase in the mean arterial blood pressure and pulse rate, and decreased the fundus pulsation amplitude, as measured by laser interferometry.^22^ Despite changes in the mean arterial blood pressure, reduced fundus pulsation amplitude may indicate a decrease in ONH deformations at a faster heart rate, which is consistent with our predicted results.

From diastole to systole, the shearing amount refers to the anterior movement of the parapapillary retina due to the choroidal expansion and the posterior ONH deformation due to both choroidal swelling and the OPA. With an increase in heart rate, it is expected that the shearing amount would decrease due to the reduced OPA and pulse volume. This shearing phenomenon has been observed in-vivo in humans using low-coherence tissue interferometry. ^18^

### The LC Became Stiffer with an Increasing Heart Rate

Our model predicted that the dynamic modulus of LC increased with an increasing heart rate. Similar phenomena have also been observed in other viscoelastic biological tissues such as the cornea^38, 39^ intervertebral discs,^40^ and other biomaterials.^41^ The dynamic modulus characterizes the stiffness of a material in response to dynamic loading, and it can differ from its static behaviour since viscoelastic materials typically become more rigid with a faster loading rate.^38, 39^ The static modulus describes the elasticity of a tissue at ‘equilibrium’, whereas the dynamic modulus provides information about the instant stiffness of a material. For instance, for the cornea, the dynamic properties are likely dominated by the rearrangement of the extracellular matrix (mainly through diffusion of water) upon changes in stress, whereas the static properties are more closely related to the collagen structure.^42^ Histological studies have shown that the morphology of the LC, as a porous, reticulated and multi-lamellar sieve-like structure, mainly comprises of laminar beams consisting of elastin fibres and collagen fibrils (type I and III).^43^ Hence, it is likely that the dynamic and static responses of the LC are also dominated by different components, and both may change differently during the progression and development of glaucoma.

### Clinical Relevance to Glaucoma Pathogenesis

At first view, a faster heart rate may be beneficial for the eye. First, it resulted in a smaller OPA and hence smaller ONH deformations. A larger deformation would yield larger stain/stress fluctuations within the LC, which may have an impact on the retinal ganglion cell axons and the vascular capillaries. This is also consistent with the fact that a larger OPA has been observed in chronic angle-closure glaucoma and suspected open-angle glaucoma subjects.^44^

Second, a faster heart rate also resulted in lower neural tissue shear, which may limit axonal damage in the neuroretinal rim region.^45-47^ As discussed in our previous work, repeated shearing of axons in the neuroretinal rim region could potentially contribute to axonal damage. The neuroretinal rim is indeed a region that exhibits a mechanical discontinuity: it is where RGC axons follow a sharp turn to enter the disc. In this region, a reduction in Bruch’s-membrane-opening minimum-rim-width (BMO-MRW) has been observed clinically in glaucoma and in elderly individuals.^45-50^ Hence, decreasing shearing with a faster heart rate may be seen as ‘mechanically protective’.

Third, the LC was found to stiffen with an increasing heart rate, which could provide better support to the RGCs and vasculatures. Pseudoexfoliation (PEX) syndrome is a systemic disorder of the elastic fiber system which can lead to PEX glaucoma. It has been reported that the stiffness of the LC in PEX eyes is 40% less than that in normal eyes as evaluated by atomic force microscopy.^51^ This decrease in LC stiffness may increase susceptibility to IOP-induced glaucomatous optic nerve damage as observed in PEX eyes, and thus, dynamic stiffening of the LC may also be seen as beneficial.

However, the situation may be more complex as the effects of dynamic loading at the cellular level has remained elusive yet. The response of cells to deformation is dependent on both the magnitude and the rate of mechanical strain. Some studies have shown that short-term strain or stress variations can lead to cellular injury in neurons and neuron-like cells.^52-54^ Future research is required to better understand the link between dynamic loading and RGC axonal injury at the micro or cellular scale.

### Understanding The Clinical Implications of LC Stiffening

During the development and progression of glaucoma, the connective tissues of the ONH undergo constant remodelling (including stiffening in the moderate/severe stages) and extensive structural changes.^55^ The extracellular matrix remodelling are usually characterized by fibrotic changes associated with cellular and molecular events such as myofibroblast activation that could drive further tissue fibrosis and stiffening.^56^ However, as discussed herein, the LC also gets stiffer with an increased heart rate. Ex vivo strain measurement shows that the stiffness of the LC increases with age.^57^ It is believed that the change in LC stiffness due to the change in heart rate and aging involves different mechanisms. Our results only reflect the short-term impact of a raised heart rate and we do not know whether the stiffening of the lamina will become chronic or whether other homeostatic mechanisms might reverse these changes in the longer term. If an increase in dynamic LC stiffness is proved to be beneficial, it is possible to modulate heart rate in the longer term via pharmocologial means. Finally, it is important to note that, while assessing the material properties of the LC from the ocular pulse, physicians should be able to distinguish whether a change in LC stiffness is because of a different heart rate/blood pressure, or because of tissue remodelling. This information may be helpful to predict glaucoma progression.

### Limitations

Our study has several limitations. First, our model did not consider the regional variations in the scleral thickness and stiffness, which are known to exist in monkeys, canines, and humans. ^20, 58, 59^ The regional variations may affect the prediction of the OPA.

Second, the choroid was simplified and modelled as a biphasic material, consisting of a porous solid phase (connective tissues) and a fluid phase (blood). In addition, the permeability of the choroid was approximated grossly from the vascular resistance, the average diameter of the choroid and the blood viscosity. ^19^ The complex microvasculature architecture, as well as the blood flow of the choroid, cannot be fully captured by this description. For instance, the segmental distribution of PCAs^60^ and the autoregulatory capacity of the choroid ^22, 61^ were not taken into account in our model. Furthermore, the PCAs and vortex veins were not modelled explicitly but represented by prescribed fluid pressures. Future work may need to assess the permeability of the choroid experimentally and also consider more complex material models to better describe the behaviour of the pulsatile choroidal blood flow.

Third, our model did not account for the blood circulation within the LC and other pulsatile components including those from the retina, central retinal artery and the CSFP. These pulsations may affect the LC deformations as well. The pulsation of the CSFP is phase-shifted with respect to the ocular pulse, such that the CSFP peak occurs slightly before that of the IOP.^62, 63^ The phase difference between the IOP and CSFP may lead to dynamic fluctuations of the translaminar pressure gradient, resulting in different ONH deformation profiles. The contribution of these pulsatile components may be important and should be considered in future studies.

Fourth, the geometry of the eye was constructed from a single eye model and the dimensions of the ONH were taken as average measurements from the literature. However, the ocular and ONH geometries can vary widely across individuals and thus may influence the predicted ocular pulse significantly. In our model, the retina, choroid and BM were simplified as a uniform layer throughout the eye. We aim to take regional variations in tissue thickness into consideration in future work.

Fifth, we experimentally measured the viscoelastic material properties of the porcine LC in this study. It is possible that the LC exhibits different viscoelastic properties across species. In addition, the LC was simplified as an isotropic and homogeneous material; the complicated laminar microarchitecture was not taken into account. Micro-level modelling of the laminar beams predicted higher strain and stress than those predicted by macro-scale models of the ONH.^64^ The mean strain of the laminar beams varies greatly across different nonhuman primates and depends on the 3D geometry of each eye’s ONH connective tissues.^64, 65^ Models incorporating the micro-level laminar beams may be considered in the future.

Sixth, for simplicity and as a first approach, we only modelled the viscoelasticity of the LC and not that of other tissues such as the choroid and the sclera. We focused on the LC only as it is a tissue that has been observed under dynamic loading^18^ and because it plays a critical role in glaucoma as the main site of damage. More complex models will be needed in the future to better understand how scleral and choroidal viscoelasticity could interact with the LC.

Seventh, the results shown only reflect the short-term impact of a raised heart rate and we do not know whether the stiffening of the lamina will become chronic or whether other homeostatic mechanisms might reverse these changes in the longer term.

Eighth, the LC was modelled as a uniform structure. We did not take into account potential differences in the anatomy and biomechanical properties of the various layers of the LC, nor did we consider lateral movements of the LC in dependence of changes of the examined parameters and in dependence of the obliqueness of the optic nerve canal and the spatial relationship between Bruch’s membrane opening and the lamina cribrosa position. ^66, 67^

## Conclusions

In this study, we used FE modelling to study the ocular pulse and the impact of the heart rate on ONH deformations. We found that the OPA, the pulse volume the ONH deformation reduce at a faster heart rate. Our models also indicated that the LC becomes stiffer with increasing heart rate. These results will help us develop technology to assess the biomechanics of the ONH in vivo by using the ocular pulse as a natural load.

